# Comparison of visualisation tools for single-cell RNAseq data

**DOI:** 10.1101/2020.01.24.918342

**Authors:** Batuhan Çakır, Martin Prete, Ni Huang, Stijn van Dongen, Pınar Pir, Vladimir Yu. Kiselev

## Abstract

In the last decade, single cell RNAseq (scRNAseq) datasets have grown from a single cell to millions of cells. Due to its high dimensionality, the scRNAseq data contains a lot of valuable information, however, it is not always feasible to visualise and share it in a scientific report or an article publication format. Recently, a lot of interactive analysis and visualisation tools have been developed to address this issue and facilitate knowledge transfer in the scientific community. In this study, we review and compare several of the currently available analysis and visualisation tools and benchmark those that allow to visualize the scRNAseq data on the web and share it with others. To address the problem of format compatibility for most visualisation tools, we have also developed a user-friendly R package, *sceasy*, which allows users to convert their own scRNAseq datasets into a specific data format for visualisation.

## Introduction

In just a decade, the number of cells profiled in each scRNAseq experiment has increased from approximately 1,000 cells to millions of cells (Svensson et al. 2017), thanks to the advent of sequencing protocols, from well-based (Deng et al. 2014; Picelli et al. 2014; Ramsköld et al. 2012) to droplet-based (Macosko et al. 2015; Klein et al. 2015), and the ever-decreasing cost of sequencing. In parallel, many computational methods have been developed to analyse and quantify scRNAseq data (Kiselev et al. 2017; Butler et al. 2018; Kiselev et al. 2018; Lee et al. 2019; La Manno et al. 2018; Stuart et al. 2019). A typical scRNAseq analysis pipeline starts from the raw reads, which are processed to create an expression matrix, containing the expression values of every gene in every cell. Further downstream analysis is then performed where cells are clustered and the cluster-specific marker genes are identified to annotate cells with corresponding cell types. The results are then visualized using non-linear embedding methods, such as tSNE (van der Maaten and Hinton 2008) or UMAP (McInnes et al. 2018) usually in a two-dimensional space where each cell gets a pair of X-Y coordinates in a two-dimensional (2D) space defining its position on the visualisation plot. Finally, the visualisations are used to assess the obtained cell types by highlighting the cell metadata (information about cells in a given experiment, e.g. batch, donor etc.) or the expression of specific genes across the cell types. This assessment can only be performed in an interactive manner. However, when the results are shared as a report or published in a paper format (a static 2D image), it is only possible to see a snapshot of the analysis corresponding to a single gene and a single set of cell metadata. Recently the ability to analyse, visualise the data in an interactive way has attracted a lot of attention, and advances in web technologies have led to the development of multiple tools for sharing the analysis results via a web interface.

In this paper we attempt to give an overview of a number of currently available tools to help researchers choose a tool for their visualisations. We compared 13 popular interactive analysis and visualisation tools for scRNAseq data by means of their features, performance and usability. Firstly, we looked at their general characteristics and properties. Secondly, we selected those tools that provide web interface functionality and benchmarked them against each other by means of their performance on datasets of different sizes (from 5,000 to 2 million cells). We also evaluated user experience (UX) features of the tools with a web interface. Finally, since all of the tools have different input requirements, we developed an R package, *sceasy*, for flexible conversion of one data format to another.

## Results

We reviewed and compared 13 popular scRNAseq analysis and visualisation tools: ASAP (Gardeux et al. 2017), BioTuring Single Cell Browser (Bbrowser) (BioTuring n.d.), cellxgene (Chan Zuckerberg Initiative n.d.), Granatum (Zhu et al. 2017), iSEE (Rue-Albrecht et al. 2018), loom-viewer (Karolinska Institutet n.d.), Loupe Cell Browser (10X Genomics n.d.), SCope (Davie et al. 2018), scSVA (Tabaka et al. 2019), scVI (Lopez et al. 2018), Single Cell Explorer (Feng et al. 2019), SPRING (Weinreb et al. 2018) and UCSC Cell Browser (UCSC n.d.). Table 1 compares these tools in terms of cloud support, containerisation, supported input formats, and developer activity.

**Table 1.** Overview of the visualisation tools and their capabilities. *Latest GitHub commit (checked on 20/12/19). **Web Interface** corresponds to the ability of hosting and sharing a webpage with a data visualisation. **Interactivity** corresponds to the ability of exploring the data in an interactive way as opposed to static images. **Docker** indicates whether a docker image with the tool is provided by the developers. **Cloud Support** indicates whether the authors provided instructions on how to deploy their web interface on a public cloud. **Loom**, **h5ad**, **SCE** (SingleCellExperiment), **Seurat**, **csv/txt** are different standard input data formats.

|  | ASAP | Bbrowser | cellxgene | Granatum | iSEE | Loom viewer | Loupe Cell Browser | SCope | scSVA | scVI | Single Cell Explorer | SPRING | UCSC Cell Browser |
| --- | --- | --- | --- | --- | --- | --- | --- | --- | --- | --- | --- | --- | --- |
| Web Interface | ✓ |  | ✓ |  | ✓ | ✓ |  | ✓ | ✓ |  | ✓ | ✓ | ✓ |
| Interactivity | ✓ | ✓ | ✓ |  | ✓ |  | ✓ | ✓ | ✓ |  | ✓ | ✓ | ✓ |
| Docker | ✓ |  | ✓ |  | ✓ |  |  | ✓ | ✓ |  | ✓ |  |  |
| Cloud Support |  |  |  | ✓ |  |  |  | ✓ | ✓ |  |  | ✓ |  |
| Loom |  |  |  |  | ✓ | ✓ |  | ✓ | ✓ | ✓ | ✓ |  |  |
| h5ad |  | ✓ | ✓ |  |  |  |  |  | ✓ | ✓ | ✓ |  | ✓ |
| SCE |  |  |  |  | ✓ |  |  |  |  |  |  |  |  |
| Seurat |  | ✓ |  |  |  |  |  |  |  |  | ✓ |  | ✓ |
| csv/txt | ✓ | ✓ |  | ✓ |  |  |  |  | ✓ | ✓ | ✓ | ✓ | ✓ |
| Platform | Java/R | Desktop | Python | R | R | Python | Desktop | Python | R | Python | Python | Python | Python |
| Last updated* | >3 m | <1 m | <1 m | >6 m | <1 m | >3 m | >3 m | <1 m | >6 m | <1 m | >3 m | >6 m | <1 m |
| Number of contributors | 2 | ? | 19 | 3 | 7 | 5 | ? | 5 | 2 | 24 | 1 | 3 | 6 |
| Number of active developers (last 2 m) | 0 | ? | 6 | 0 | 3 | 0 | ? | 2 | 0 | 4 | 0 | 0 | 2 |
| Version | 1.0 | 2.2.13 | 0.11.0 | 0.0.0.900 | 1.4.0 | 0.32.4 | 3.1.1 | 1.7.2 | 0.2.0 | 0.5.0 | 1.0.0 | 1.6.0 | 0.5.49 |

The tools vary in the ability to use different input file/data formats (green colour in Table 1). We focused on input formats supported out of the box. The csv/txt format is the most accepted one and can be used by eight tools. More specialized formats such as h5ad and loom are accepted by six tools. R-based SingleCellExperiment (SCE) and Seurat are accepted by one and three tools, respectively. To make it possible for the users to visualize their datasets in different ways we have developed the *sceasy* R package for file format conversion, which is available on Github at https://github.com/cellgeni/sceasy.

In terms of web features (blue colour in Table 1), some of the tools are provided with cloud-specific deployment instructions (Granatum, SCope, scSVA and SPRING) whereas others (cellxgene, iSEE, SCope, scSVA, Single Cell Explorer) have Docker images that facilitate deployment, but require more development work. ASAP is a comprehensive hosting platform at École Polytechnique Fédérale de Lausanne (EPFL) and as such it does not require a local or cloud installation. Additionally it provides a Docker image for custom deployment. Note that not every web-friendly tool has the ability to host and share a webpage with a data visualisation (**Web Interface** row in Table 1). For example, Granatum provides a full analysis pipeline which includes some visualisations, however, they do not currently allow for sharing them with others.

For further comparison we selected actively developed (updated during the last six months) tools which can be used for hosting and sharing a web page with data visualisation. For ASAP we were not able to host it locally with the provided Docker image, hence it was excluded from the comparison.

For the end user to see a visualisation web page it usually requires three steps to be completed: the input files have to be prepared (e.g. to create a database or to convert to another data format, different from the ones in Table 1), the back-end server has to be started and, finally, the web page should be served to the user in their web browser. For all of the considered tools the memory and time needed for the latter step are negligible compared to the first two steps. We defined the maximum memory and total time required for the first two steps as the preprocessing memory (RAM) and the preprocessing time and measured how they depend on the number of cells in the input data. All of the tools were run with the default parameters. iSEE was run for both SCE and loom input formats.

Fig. 1 summarizes the benchmarking results. iSEE-loom, SCope, scSVA and loom-viewer all enable efficient integration with the hierarchical data format (HDF5) from which loom and h5ad formats are derived. HDF5 format allows for on-demand loading, i.e. the necessary data is only loaded in RAM when it is needed by the application. In this case, at the start the tools only use the coordinates of the cells without loading other input data into memory. scSVA and loom-viewer are the most efficient HDF5-backed tools, and SCope is slightly slower. iSEE-loom is memory efficient but there is a sudden increase in the preprocessing time at 250K and 500K cells. This is due to having 8 panels in the default iSEE interface, each requiring a separate rendering of the visualisation of the input cells. The defaults can be programmatically changed to having only one panel. This brings the preprocessing times down to under 2 minutes for 500K cells. Due to long preprocessing times we did not run iSEE-loom for datasets with more than 500K. Similarly, there is a sudden drop of loom-viewer efficiency at 2M cells. This effect was consistent across all 5 runs but we could not explain it. There is also a consistent drop in RAM usage by Single Cell Explorer at 50K cells, which we could not explain. In addition, we were not able to run SCope and Single Cell Explorer for datasets larger than 500K cells.

**Figure 1.**
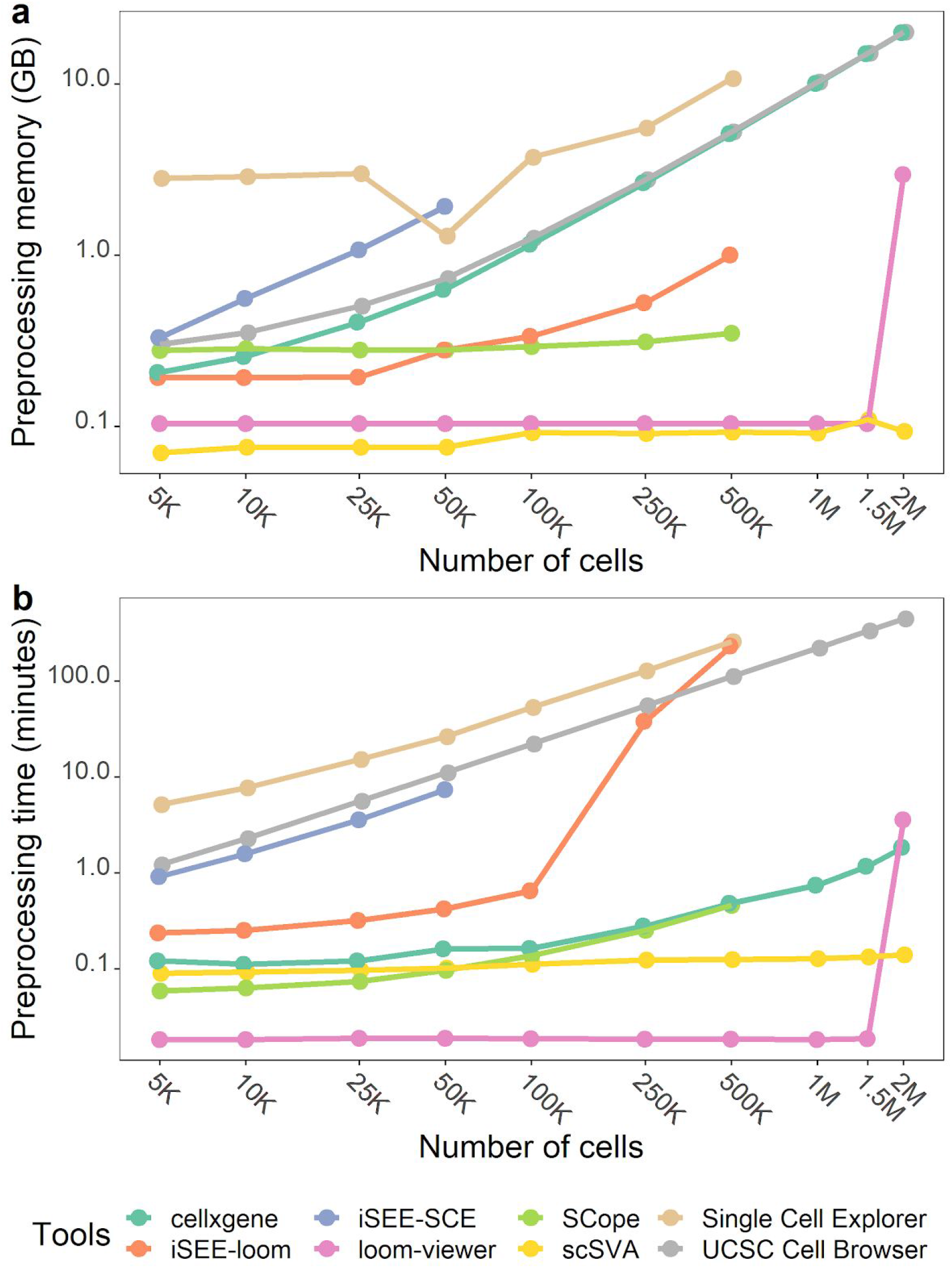
Preprocessing RAM usage (a) and preprocessing times (b) of the visualisation tools. The points on the plots represent median times across 5 independent runs. Preprocessing times include input preparation and starting of the back-end server.

For four tools (iSEE-SCE, Single Cell Explorer, UCSC Cell Browser and cellxgene) the preprocessing memory and preprocessing time grow exponentially with the number of input cells. In case of iSEE-SCE and cellxgene this is due to the lack of HDF5 integration and loading of the full data in memory, which increases the server starting time. In contrast, the long preprocessing times of Single Cell Explorer and UCSC Cell Browser are explained by the required database and file preparation time, respectively. iSEE-SCE has the steepest RAM usage growth with the number of cells and failed to start for datasets larger than 50K cells. Among these four tools cellxgene has the shortest preprocessing times.

In addition to the performance we also scored the benchmarked tools by their user experience as shown in Table 2.

**Table 2.** UX scores of the visualisation tools.

|  | cellxgene | iSEE | Loom-viewer | scSVA | SCoPe | Single Cell Explorer | UCSC Cell Browser |
| --- | --- | --- | --- | --- | --- | --- | --- |
| Ease of cell selection | ✓ | ✓ |  | ✓ | ✓ | ✓ | ✓ |
| Zoom in/out | ✓ | ✓ |  | ✓ | ✓ |  | ✓ |
| Multiple embeddings | ✓ | ✓ | ✓ |  | ✓ |  | ✓ |
| Highlight gene expression | ✓ | ✓ |  | ✓ | ✓ | ✓ | ✓ |
| Highlight metadata | ✓ | ✓ | ✓ | ✓ |  | ✓ | ✓ |
| Extra analysis | ✓ | ✓ |  | ✓ |  | ✓ |  |
| Web page loads fast | ✓ |  | ✓ | ✓ | ✓ | ✓ | ✓ |

For statistical analysis and comparison of groups of cells users need to select cell populations of interest. A majority of the tools provide selection functionality. The most flexible and user-friendly of them is the free-hand lasso selection, which is supported by cellxgene, SCope and Single Cell Explorer. It allows the user to select the cells by drawing a free shape curve around the cells of interest. A less flexible type of selection is rectangular selection, where the user is limited to drawing a rectangle around the cells of interest. It is supported by scSVA and UCSC Cell Browser. In addition, iSEE also supports a polygon version of the lasso selection, where the selection is made by pointing to the vertices of a polygon. To our knowledge, loom-viewer does not provide any method of selection.

The ability to zoom in and out can be crucial to visually analyse and validate the data. Most of the tools have zooming functionality except loom-viewer and Single Cell Explorer. Similarly, the ability to switch between multiple embeddings (e.g. between tSNE and UMAP) can be very useful and help with the analysis. Again most of the tools have this functionality, except scSVA and Single Cell Explorer.

One of the most important features every single-cell visualisation tool must have is the capability to highlight specific information. The user may want to highlight either gene expression levels (continuous scale) or cell metadata (usually on a discrete scale). Not surprisingly, this functionality is available in almost every tool with the exception of loom-viewer (for gene expression) and SCope (for cell metadata).

A useful feature of a visualisation tool is the option of performing extra analysis on user-selected cells, such as e.g. cell-type annotation, differential expression analysis or marker gene identification. Three of the benchmarked tools (cellxgene, scSVA and Single Cell Explorer) have this functionality out of the box. iSEE supports this via custom panels.

Finally, an important indicator of a good user experience is the speed of web page loading. Once the back-end server is running most of the tools can serve the visualisation to the user via a web page in a fast manner. One exception is iSEE. We tested it with the default setting of 8 panels. At the moment the free version of the Shiny server does not support persistent R processes for faster load times (https://rstudio.com/products/shiny/shiny-server/) and therefore it starts a new R process for each new user, increasing the loading times and affecting the user experience when data sets are large.

## Methods

### Datasets

For benchmarking we utilised a mouse embryo development scRNAseq dataset with an accession number of GSE119945 (Cao et al. 2019) which contains 2.07 million cells. Ten datasets of different sizes (with 5,000, 10,000, 25,000, 50,000, 100,000, 250,000, 500,000, 1,000,000, 1,500,000 and 2,000,000 cells) used for performance benchmarking were created by randomly subsampling cells of the original dataset.

### Profiling

Benchmarking tests were done on a virtual Ubuntu OS 16.04 with 23GB of RAM and 2GHz Intel Xeon Processor with 16 cores.

iSEE and scSVA are both R packages and therefore were tested by using *profvis*, a package for profiling R scripts (Chang et al. 2019). The highest value of the *“memalloc”* slot with the label of *“shiny::runApp”* was considered as preprocessing memory, and the last value in the *“time”* slot was considered as preprocessing time.

For all the other tools (except cellxgene) the characteristics were measured by running Linux command `/usr/bin/time -v`, and using *“Maximum resident set size (kbytes)”* output for RAM usage and using the sum of *“User time (seconds)”* and *“System time (seconds)”* outputs for preprocessing times. For cellxgene the *gnomon* command was used with elapsed-total option to measure the preprocessing times. For SCope the preprocessing time was profiled only from the server side (the timed process was `scope-server`). The preprocessing times included both the internal data import time (only for UCSC Cell Browser and Single Cell Explorer) and server start-up time. Both UCSC Cell Browser and Single Cell Explorer spent a considerable amount of time on the data import as they need to convert the data into other structures (json files and MongoDB database, respectively) before being able to visualize it. For UCSC Cell Browser, scanpy’s (Wolf et al. 2018) `scanpy.external.exporting.cellbrowser` was used to perform the conversion. For Single Cell Explorer, `ProcessPipline.insertToDB` function from the `scpipline.py` library provided by the authors was used.

## Discussion

The size and volume of scRNAseq data has exponentially increased over the last decade and this has opened up new avenues of scientific discovery and understanding. There is a need among scientists to communicate their data to collaborators and colleagues for quick and easy exploration. The burden of computational resources and bioinformatics skills required to do this should ideally be removed from the recipients of the data.

Single-cell interactive analysis and visualisation tools have been widely adopted by the research community. They make data import, public data access and analysis much easier for the users and accelerate the science. Furthermore, tools now exist (those with **Web Interface** functionality in Table 1) that allow the user to host and share their scRNAseq data visualisation with others on the web, due to recent advances in web technologies. These make it possible to share analysis results with others in a user-friendly manner, allowing for much faster scientific development. We believe that high complexity and dimensionality of scRNAseq data can only be revealed via comprehensive, interactive, and user-friendly tools that can provide shareable visualisations via the web. This is supported by the recent developments of scRNAseq visualisation portals at large scientific institutes:

1. Single-Cell Expression Atlas (Papatheodorou et al. 2020) at the European Bioinformatics Institute - https://www.ebi.ac.uk/gxa/sc/home
2. Single-Cell Portal at the Broad Institute - https://singlecell.broadinstitute.org/single_cell
3. Cell Browser at the University of California Santa Cruz - https://cells.ucsc.edu/
4. Automated Single-cell Analysis Pipeline (ASAP) at École Polytechnique Fédérale de Lausanne - https://asap.epfl.ch/

To understand the current landscape of interactive analysis and visualisation tools we compared (Table 1) several of the most popular based on their general qualities. Our results show that each tool has particular advantages and disadvantages and as such a simple ranking cannot be achieved. We looked specifically at the tools suitable for sharing an interactive visualisation of results via a web interface (the **Web Interface** row in Table 1). Again, in this case there is not one tool that stands out as significantly better than the rest in all categories. From our personal experience and partly supported by benchmarking, we currently recommend using cellxgene for publishing and sharing scRNAseq data.

Cellxgene performs well in terms of both memory and preprocessing times (Fig. 1) and leads by the UX scores (Table 2). It also has a thriving community, is the most supported (Table 1) with 24 contributors and has the highest developer activity in the last 2 months. We have developed a detailed tutorial (https://cellgeni.readthedocs.io/en/latest/visualisations.html) for using cellxgene and how to convert data into the required input format.

Single-cell sequencing technologies are still in rapid development and we expect the dataset sizes (number of cells per dataset) to further grow in the next few years. Tools that use on-demand loading with linear or sublinear memory usage relative to cell count are best positioned to cope with this growth. Other tools will have to adapt and optimise in order to stay competitive. One way of optimisation is to add integration with the HDF5 format as supported by our results in Fig. 1. A simpler approach is to down-sample the data by selecting a small subset of cells in a random manner. Most of the tools support this functionality. When down-sampling it is important to make sure that rare cell populations are not removed. One example approach is geometric sketching (Hie et al. 2019) - a method to subsample massive scRNA-seq datasets while preserving rare cell states.

Additionally, there are several efforts to enrich visualisations by either using a 3D plot instead of 2D (Hillje et al. 2019; Gardeux et al. 2017) or even by using virtual reality (Legetth et al. 2018). The third dimension may allow resolution of cell-types not visible in two dimensions. Virtual reality can support multiple embeddings in the same VR space so that they can be directly compared.

This review represents a snapshot of a rapidly developing field and tools will catch up or drop out of contention and new tools will emerge. All of the tools in this review that are under active development are worth keeping an eye on. The designers and developers will need to not only think about efficiency and scalability of visualisation but also about additional features that can enrich the data visualisation and provide more scientific insights. An example is the user-friendly integration of a dataset under consideration with public scRNAseq data. This is already happening in commercial products, e.g. Bbrowser provides, for a selected group of cells, a suggestion of cell type based on publicly available data. It also provides the ability to search for specific cells from public data similar to the selected ones. It is worth noting that command line tools exist with exactly the same or similar functionality. However, putting this functionality into an interactive user-friendly interface allows sharing and exploration of results across the whole research community, facilitating scientific progress.

## Acknowledgements

We would like to thank Maximilian Haeussler and Shamit Soneji for their constructive comments and feedback.

